# Associations between aversive learning processes and transdiagnostic psychiatric symptoms revealed by large-scale phenotyping

**DOI:** 10.1101/843045

**Authors:** Toby Wise, Raymond J Dolan

## Abstract

**Background:** Symptom expression in a range of psychiatric conditions is linked to altered threat perception, manifesting particularly in uncertain environments. How precise computational mechanisms that support aversive learning, and uncertainty estimation, relate to the presence of specific psychiatric symptoms remains undetermined. 400 subjects completed an online game-based aversive learning task, requiring avoidance of negative outcomes, in conjunction with completing measures of common psychiatric symptoms. We used a probabilistic computational model to measure distinct processes involved in learning, in addition to inferred estimates of safety likelihood and uncertainty, and tested for associations between these variables and traditional psychiatric constructs alongside transdiagnostic dimensions. We used partial least squares regression to identify components of psychopathology grounded in both aversive learning behaviour and symptom self-report. We show that state anxiety and a transdiagnostic compulsivity-related factor are associated with enhanced learning from safety, and data-driven analysis indicated the presence of two separable components across our behavioural and questionnaire data: one linked enhanced safety learning and lower estimated uncertainty to physiological anxiety, compulsivity, and impulsivity; the other linked enhanced threat learning, and heightened uncertainty estimation, to symptoms of depression and social anxiety. Our findings implicate distinct aversive learning processes in the expression of psychiatric symptoms that transcend traditional diagnostic boundaries.

## Introduction

Many core symptoms of mental illness are linked to learning about unpleasant events in our environment. In particular, symptoms of mood and anxiety disorders, such as apprehension, worry, and low mood can intuitively be related to altered perception of the likelihood of aversive outcomes. Indeed, the importance of altered threat perception is a feature of many diagnoses that extend beyond disorders of mood to encompass conditions such as psychosis^1^ and eating disorders^2^. As a result, research into how individuals learn about aversive events holds great promise for enhancing our understanding across a diverse range of mental health problems.

Computational approaches are a powerful means to characterise the precise mechanisms underpinning learning, as well as uncovering how these relate to psychiatric symptom expression^3,4^. Recent studies have leveraged computational modelling to capture associations between learning processes and psychiatrically-relevant dimensions in non-clinical samples^5–8^, as well as in clinical conditions ranging from anxiety and depression to psychosis^9–12^. A common finding across studies is that of altered learning rates, where psychopathology is linked to inappropriate weighting of evidence when updating value estimates^7,13,14^. Notably, there is evidence suggesting that people with clinically significant symptoms of anxiety and depression show biased learning as a function of the valence of information, updating faster in response to negative than positive outcomes presented as monetary losses and gains^12^, a bias that might engender a negative view of the environment. However, we previously found an opposite pattern in a non-clinical study using mild electric shocks as aversive stimuli, whereby more anxious individuals learned faster from safety than from punishment, and underestimated the likelihood of aversive outcomes^15^. This latter finding highlights a need for a more extensive investigation using larger samples.

In addition to aberrant learning another process implicated in the genesis of psychiatric disorder relates to the estimation of uncertainty^16^. While there are multiple types of uncertainty, here we use the term to refer to estimation uncertainty, describing the precision of a learned association. Estimation uncertainty is highest when there is a lack of experience, or the association to be learned is unstable. For example, having seen two coin flips and observing one head and one tail, one might believe the likelihood of observing a head is 50%, though you are highly uncertain about this estimate due to a lack of evidence. This kind of uncertainty plays a fundamental role in learning, and computational formulations optimise learning in the face of non-stationary probabilistic outcomes based on uncertainty^11,17–20^. While psychiatric symptoms, including anxiety, have been linked to an inability to adapt learning in response to environmental statistics such as volatility^5,9^, little research has investigated how individuals estimate, or respond to, uncertainty in aversive environments and its potential association with psychiatric symptoms. This is a crucial question given that core features of anxiety revolve around a concept of uncertainty. For example, individuals with anxiety disorders report feeling more uncertain about threat and being less comfortable in situations involving uncertainty^21–24^. Surprisingly, in an earlier lab-based study we observed a surprising relationship, finding that more anxious individuals were more certain about stimulus-outcome relationships^15^. However, this was in a relatively small sample and therefore warrants further investigation.

Existing work on aversive learning has had a particular focus on symptoms of anxiety and depression^7,12^. However, these approaches have not been designed optimally for identifying mechanisms that span traditional diagnostic boundaries. This assumes importance in light of recent studies, using large samples, showing several aspects of learning and decision making relate more strongly to transdiagnostic factors (symptom dimensions that are not unique to any one disorder) than to any specific categorical conception of psychiatric disorder^6,8,25–27^. Applying such an approach to aversive learning may yield better insights into the role of learning in psychiatric disorders. Additionally, computationally-defined measures of learning and decision making can facilitate identification of novel transdiagnostic factors, going beyond those identified based solely on correlated symptom clusters in self-report and clinical interview measures^6,28–30^.

Here, we aimed to clarify the nature of the relationship between aversive learning processes and traditional measures of anxiety, as well as transdiagnostic psychiatric factors identified in prior work^6^ in a large, preregistered study conducted online. This allowed us to measure effects with high precision, potentially helping to resolve mixed findings from previous studies^12,15^, in addition to identifying small but meaningful effects that cross traditional diagnostic boundaries^6^. Thus, we used a computational approach to test whether anxiety and transdiagnostic symptoms are associated with biased learning from safety and threat, whether these factors relate to altered estimates of threat likelihood, and whether they are associated with different levels of uncertainty during threat learning. We then used partial least squares (PLS) regression, a data-driven multivariate method, to derive transdiagnostic latent components of psychopathology grounded in both self-report and computational measures. Given difficulties in using traditional aversive stimuli in an online setting, we developed a game-based avoidance task designed to engage threat and avoidance processes without the need for administration of painful or noxious stimuli. Both the task and modelling are, in principle, similar to our previous lab-based task^15^, but their implementation here allows straightforward administration in large samples recruited online.

## Results

### Task performance

Four hundred subjects recruited online through Prolific^31^ performed a game-based aversive learning task, where the aim was to fly a spaceship through asteroid belts without being hit (Figure 1). Getting hit by the asteroids reduced the integrity of the spaceship, and after sufficient hits the game terminated. Crucially, there were two zones at the top and bottom of the screen where subjects could encounter a hole in the asteroid belt, each associated with a changing probability of being safe. In order to perform well at the task subjects needed to learn which zone was safest and behave accordingly.

**Figure 1.**
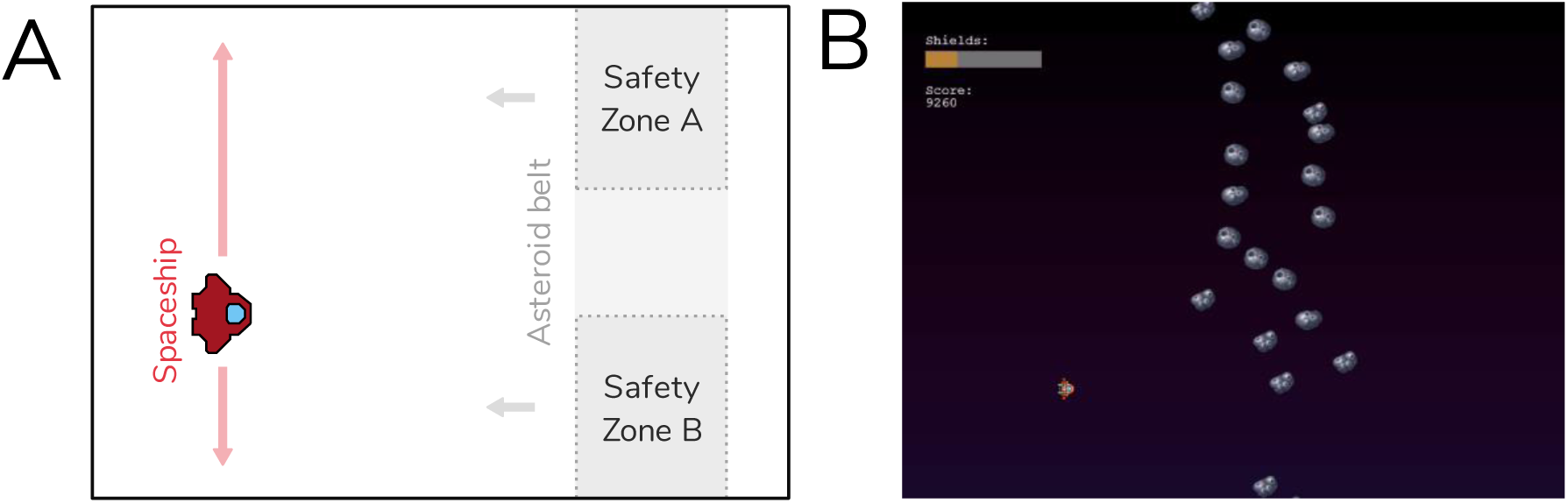
A) Task design. Subjects were tasked with playing a game that had a cover story involving flying a spaceship through asteroid belts. Each asteroid belt featured two locations that could potentially contain escape holes (safety zones), and subjects were instructed to aim to fly their spaceship through these to gain the highest number of points. Subjects were only able to move the spaceship in the Y-dimension, while asteroid belts moved towards the spaceship. The probability of each zone being safe varied over the course of the task but this could be learned, and learning this probability facilitated performance. B) Screenshot of the task, showing the spaceship, an asteroid belt with a hole in the lower safety zone (safety zone B), a representation of the spaceship’s integrity (shown by the coloured bar in the top left corner) and the current score.

Subjects were engaged and performed well at the task, with a median number of spaceship destructions of 1 (Interquartile range = 2) over the course of the task. They also reported high motivation to perform the task, providing a mean rating of 85.70 (SD = 18.44) when asked to rate how motivated they were to avoid asteroids on a scale from 0-100.

### Computational modelling of behaviour

To quantitatively describe behaviour, we fit a series of computational models to subjects’ position data during the task (see Methods and Supplementary Material for a full description of tested models). The winning model was a probabilistic model incorporating different updates parameters for safety and danger, as well as a “stickiness” parameter representing a tendency for subjects to stick with their previous position. This model represents an extension of one we have previously used successfully as a lab-based aversive learning task^15^, and is described fully in the Methods section. Briefly, this winning model assumes that subjects in the task represent the safety probability of each zone using a beta distribution, which is updated on each trial based on encounters with danger or safety. Simulating responses using the model, using each subject’s estimated parameter values, produced behavioural profiles that demonstrated a high concordance with the true data, reproducing broad behavioural patterns seen in the true data (Figure S1).

### Task and model validation

In the light of the task’s novelty, it was important to ensure the task has content validity, and that it produces behaviour reminiscent of more traditional tasks. Likewise, the computational models used should provide measures and parameter estimates that reflect the behaviour they aim to describe. We therefore conducted extensive validation exercises. These are reported fully in supplementary material, but we summarise these here.

First, we ensured the task did induce states of subjective anxiety in the majority of subjects (Figure 2A), and this level of anxiety was correlated with self-report state and trait anxiety (Figure 2C and 2D). Importantly, the task produced behaviour reminiscent of more traditional lab-based tasks, with subjects adjusting their position to a greater extent following danger than following safety (Figure 2E), as demonstrated in prior studies^12,15,32^. With respect to our computational model, we verified the model’s update parameters were robustly correlated with subjects’ tendency to move, or stay, following danger and safety respectively (Figure 3D). We also ensured that the safety value and uncertainty values produced by simulating data from our model with best fitting parameters correlated with subjects’ model-free behaviour, finding that subjects changed their position more when model-derived uncertainty was high, and when the difference between the safety value of the two zones was small, as expected (Figure 3C). Finally, we verified that our model’s update parameters showed greater updating from danger relative to safety, as we found in a previous lab-based study^15^, finding this was indeed the case (Figure 3E).

**Figure 2.**
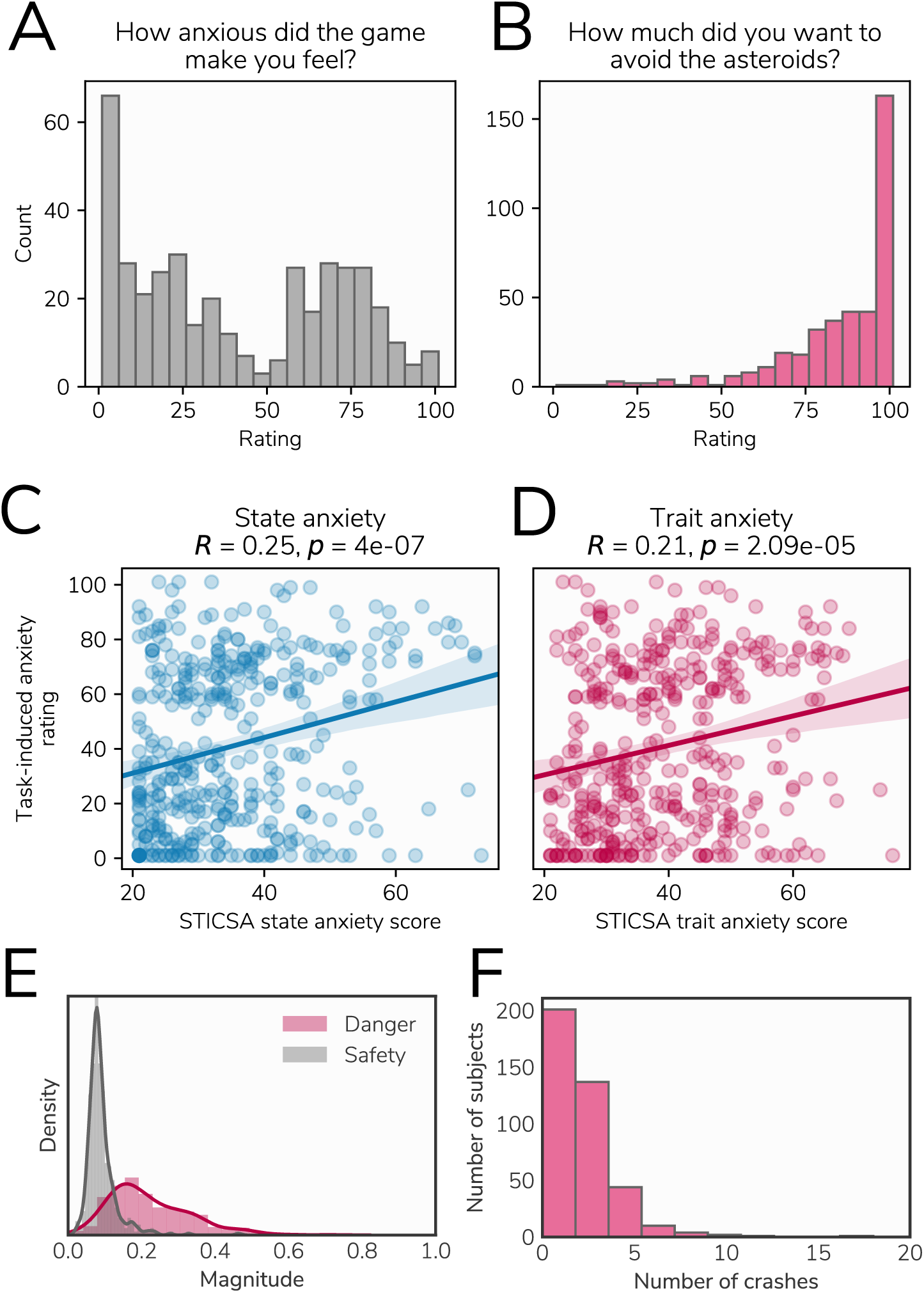
A) Distribution of task-induced anxiety ratings recorded after the task. B) Distribution of task motivation ratings. C and D) Relationships between task-induced anxiety ratings and state and trait anxiety scores. E) Degree of location switching after encountering danger and safety across subjects. The switch magnitude is the average absolute change in position between trial *n* and trial *n+1*. As expected, subjects showed more switching behaviour after encountering danger and were more likely to stay in the same position following a safe outcome. F) Distribution of crash number (representing the number of subjects hit enough asteroids to end the game) across subjects.

**Figure 3.**
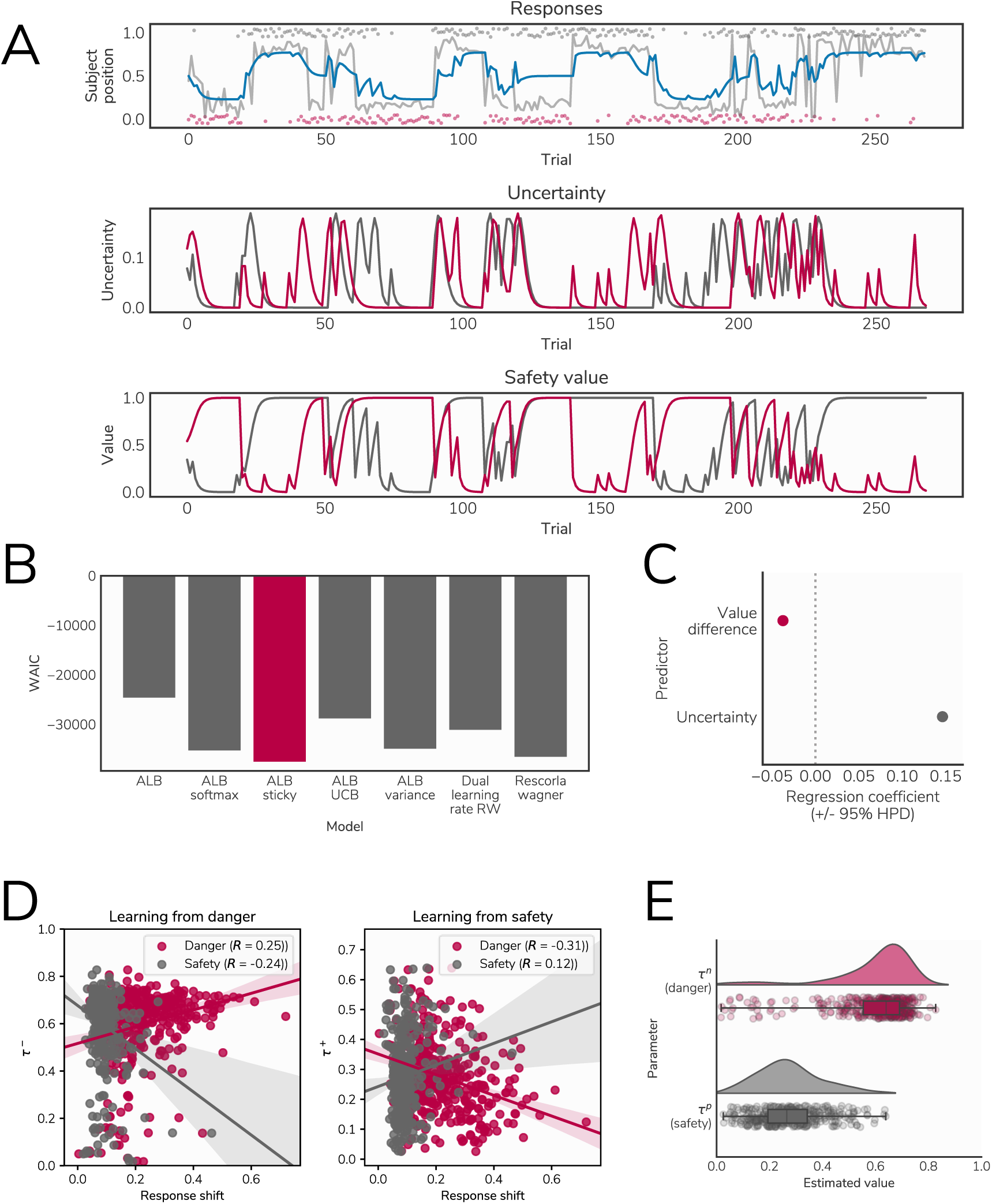
A) Data generated from the model. The top panel shows responses and model fit for an example subject. In the top panel, the grey line represents the subject’s position throughout the task, with the grey and red dots representing safe locations on each trial. The blue line represents simulated data from the model for this subject. The lower two panels show estimated uncertainty and safety probability for each stimulus (represented by the grey and red lines) across the duration of the task, generated by simulating data from the model. B) Model comparison results, showing the WAIC score for each model with the winning model highlighted. ALB = asymmetric leaky beta, RW = Rescorla-Wagner. C) Results of our analysis validating the safety value and uncertainty measures, showing the extent to which each measure predicted subjects’ tendency to switch position (described in supplementary materials). D) Correlations between estimated update parameters for danger (left) and safety (right) and our behavioural measure of position switching after these outcomes across subjects, demonstrating that parameters from our model reflect purely behavioural characteristics. E) Distributions of estimated parameter values for τ^+^ and τ^-^, representing update rates following danger and safety outcomes respectively, showing a bias in updating whereby subjects update to a greater extent in response to danger than safety.

### Relationships with psychiatric measures

First, we asked whether our four behavioural variables of interest (threat update parameter, danger update parameter, mean estimated safety probability, and mean estimated uncertainty) were associated with anxiety (both state and trait) and intolerance of uncertainty. The strongest relationships, with HPD intervals that did not include zero, were positive effects of state anxiety on safety update rates and mean estimated safety probability (Figure 4, Table 1), although effects for trait anxiety were in the same direction and of a similar magnitude for some measures, indicating more anxious individuals learned faster about safety and perceived safety as more likely overall.

**Table 1.** Estimates from regression model predicting learning-related variables derived from our computational model from measures of intolerance of uncertainty, state anxiety, and trait anxiety. Effects with HPDIs excluding zero are shown in bold. IUS: Intolerance of uncertainty scale; STICSA S: State-trait inventory of cognitive and somatic anxiety, state measure; STICSA T: State-trait inventory of cognitive and somatic anxiety, trait measure; HPDI: Highest posterior density estimate

| Target variable | Predictor | Estimate (+/- 95% HPDI) |
| --- | --- | --- |
| Threat update ($\tau^-$) | IUS | 0.07 (-0.03, 0.16) |
|  | STICSA S | -0.09 (-0.19, 0.01) |
|  | STICSA T | -0.03 (-0.12, 0.07) |
| Safety update ($\tau^+$) | IUS | 0.0 (-0.1, 0.09) |
|  | STICSA S | **0.12 (0.02, 0.21)** |
|  | STICSA T | 0.09 (-0.01, 0.18) |
| Mean safety uncertainty | IUS | -0.05 (-0.14, 0.05) |
|  | STICSA S | -0.09 (-0.19, 0.01) |
|  | STICSA T | -0.08 (-0.18, 0.01) |
| Mean safety probability | IUS | -0.02 (-0.12, 0.07) |
|  | STICSA S | **0.12 (0.02, 0.21)** |
|  | STICSA T | 0.06 (-0.03, 0.15) |

**Figure 4.**
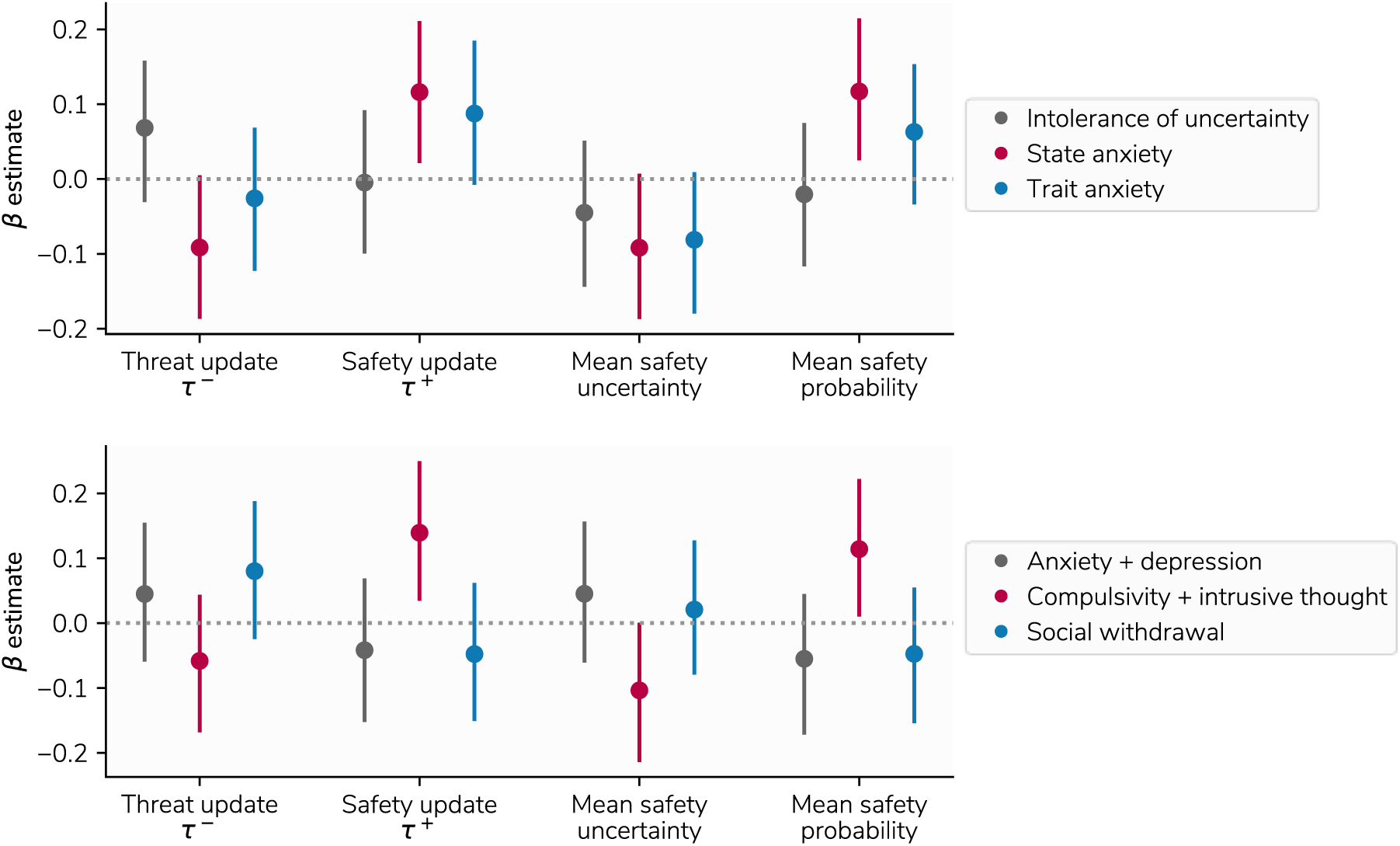
Top panel: Results of state/trait anxiety and intolerance of uncertainty models, showing relationships between these psychiatric variables and behavioural variables. Points indicate the mean of the posterior distribution for the regression coefficient parameter, while error bars represent the 95% highest posterior density interval. The *β* estimate here refers to the regression coefficient for each predictor. Bottom panel: Results of three factor model, showing relationships between behaviour and factors labelled anxiety and depression, compulsivity and intrusive thought, and social withdrawal.

We then examined the extent to which task behaviour was associated with three transdiagnostic factors of psychopathology identified through self-report assessments in previous research^6^. Here, we observed effects of a factor labelled compulsivity and intrusive thought (Figure 4, Table 2), reflecting the fact that subjects scoring higher on this factor learned faster about safety and had higher safety probability estimates. There was also a weak effect of this factor on uncertainty, although the HPDI for this included zero. Other effects were weak, and including reported task motivation as a covariate had a negligible effect on the results (see supplementary results). Importantly, all of these analyses were determined *a priori* and are included in our preregistration.

**Table 2.** Estimates from regression model predicting learning-related variables derived from our computational model from the three transdiagnostic factors identified by Gillan et al. (2016). Effects with HPDIs excluding zero are highlighted. AD: Anxious-depression; CBIT: Compulsive behaviour and intrusive thought; SW: Social withdrawal; HPDI: Highest posterior density estimate

| Target variable | Predictor | Estimate (+/- 95% HPDI) |
| --- | --- | --- |
| Threat update ($\tau^-$) | AD | 0.04 (-0.06, 0.15) |
|  | CBIT | -0.06 (-0.17, 0.04) |
|  | SW | 0.08 (-0.02, 0.19) |
| Safety update ($\tau^+$) | AD | -0.04 (-0.15, 0.07) |
|  | CBIT | **0.14 (0.03, 0.25)** |
|  | SW | -0.05 (-0.15, 0.06) |
| Mean safety uncertainty | AD | 0.05 (-0.06, 0.16) |
|  | CBIT | -0.1 (-0.21, 0.0) |
|  | SW | 0.02 (-0.08, 0.13) |
| Mean safety probability | AD | -0.06 (-0.17, 0.04) |
|  | CBIT | **0.11 (0.01, 0.22)** |
|  | SW | -0.05 (-0.15, 0.05) |

### Psychiatric constructs derived from behaviour and self-report

Numerous studies have used dimensionality reduction procedures such as factor analysis on questionnaire-based data to identify factors of psychopathology that cut across diagnostic boundaries^6,28–30^. This, in turn, has revealed that many behaviourally-defined phenotypes are more strongly associated with transdiagnostic factors than any single disorder^6,8,26^. We built upon this work by incorporating computationally-derived indexes of behaviour into this dimensionality reduction procedure, where the aim was to identify latent constructs grounded in both self-report and behaviour. We used partial least squares (PLS) regression, a method that identifies latent components linking multivariate data from multiple domains based on their shared covariance. This method has been employed successfully to provide insight into how panels of cognitive and behavioural measures relate to multivariate neuroimaging-derived phenotypes^33–35^. We first identified the number of components that best describe our data by evaluating the performance of a predictive PLS model using cross-validation. We found two latent components gave the best predictive performance (Figure 5A). We then evaluated the performance of this model on held out data using permutation testing, showing our model achieved a statistically significant level of predictive accuracy (permutation *p* = 0.025, Figure 5B). This indicates that our combined self-report and behavioural data is best explained by a two-component structure linking these two domains, Importantly, the fact that this level of accuracy was found on unseen data ensures that our results do not result from overfitting the training data^36^.

**Figure 5.**
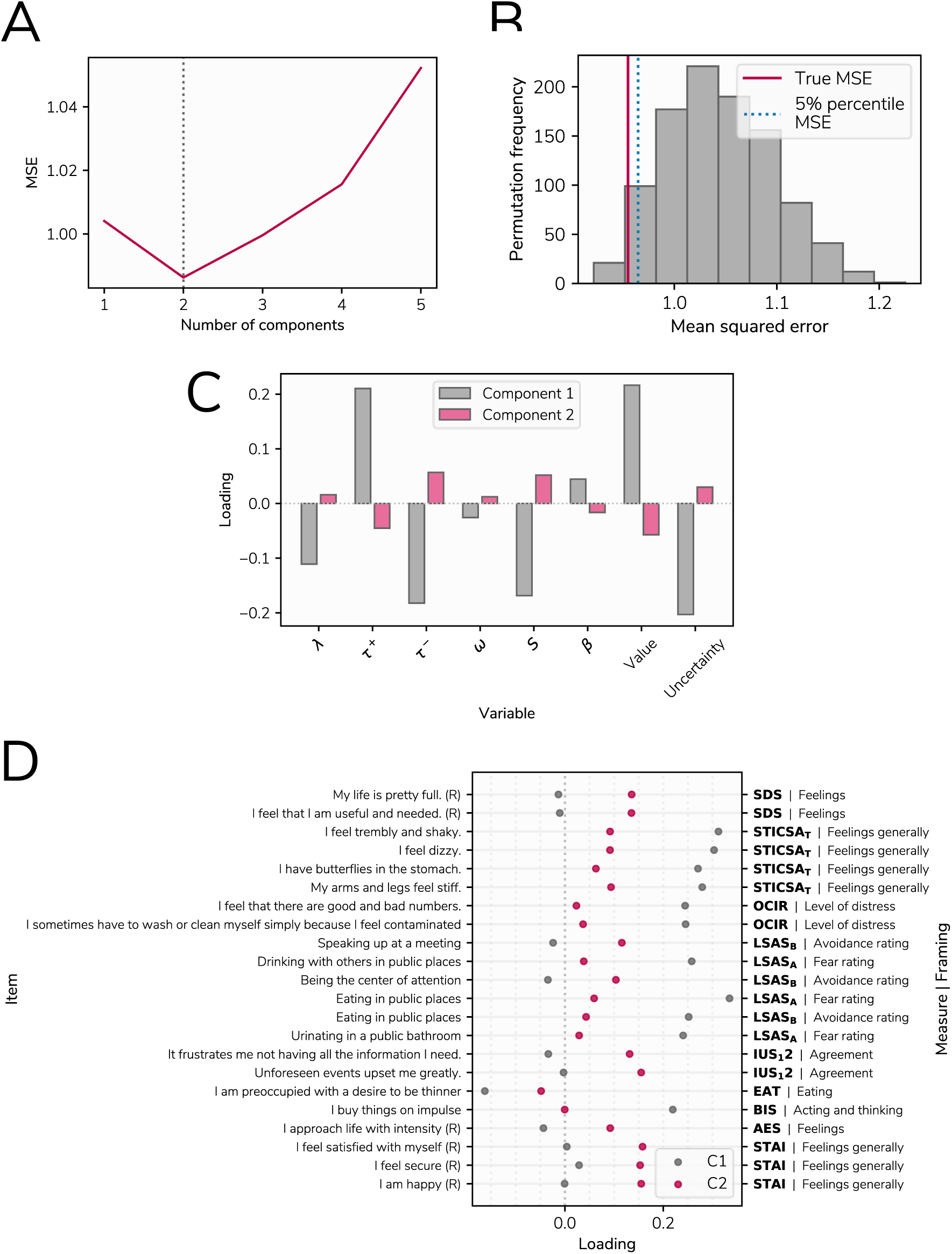
Results of PLS regression analysis. A) The optimal number of components was determined based on cross-validated predictive accuracy within 75% of the data used for training the model. This figure represents the negative mean squared error of these predictions across models with between one and five factors, showing best performance with 2 components. B) Null distribution of predictive accuracy scores generated by retraining our PLS regression model on 1000 permuted datasets and testing on the held out 25% of the data set, with the mean squared error (MSE) achieved by the model trained on the true data shown by the red line. C) Loadings for the two components on behavioural variables, including all parameters in the model and mean safety probably and uncertainty estimates, across all subjects. D) Loadings on questionnaire items showing the largest dissociations in loadings between the two components, identified by taking the lowest and highest 10% of differences between loadings. Items marked (R) are reverse coded. The labels on the right indicate the measure the item is taken from and an indicator of how the question is framed. SDS: Zhung Self-Rating Depression Scale, STICSA: State Trait Inventory of Cognitive and Somatic Anxiety, OCIR: Obsessive Compulsive Inventory, LSAS: Liebowitz Social Anxiety Scale (A and B represent subscales), IUS12: Intolerance of Uncertainty Scale, EAT: Eating Attitudes Test, AES: Apathy Evaluation Scale, STAI: State Trait Anxiety Inventory.

To aid interpretation of these two components we examined how behavioural variables loaded on each. The first component had positive weights on update rates in response to safety and estimated safety likelihood, and negative weights on update rates in response to threat, decay, stickiness, and mean uncertainty estimates (Figure 5C), while the reverse was true of the second component. Loadings on questionnaire items were varied, and labelling such components is invariably subjective. Nevertheless, the first component tended to load most strongly on items describing physical symptoms of anxiety, compulsive behaviour, and impulsivity. In contrast, the second latent component loaded primary on items describing social anxiety and depressed mood. For illustrative purposes, items with the top 10% percent of differences in loadings between components are shown in Figure 5D, with full details available in supplementary material.

## Discussion

Perceptions of danger and safety have been linked to key symptoms of psychiatric disorders. Here, in a large-scale study examining aversive learning we show that when subjects learn to avoiding threat, transdiagnostic components of psychopathology relate to how they learn about both safety likelihood, and uncertainty.

We found a counter-intuitive relationship between biases in learning and the presence of features of anxiety. Subjects scoring higher on state anxiety tended to update their predictions to a greater extent in response to safety, as well as perceiving safety to be more likely overall, than those scoring low on this measure. These results diverge from previous findings that report individuals diagnosed with clinical anxiety and depression learn faster from punishment^12^, but are in concordance with our previous work in a non-clinical sample using a more traditional lab-based aversive learning task^15^. The large sample size employed here allowed us to estimate these effects precisely, making it unlikely that they are simply a product of statistical noise. One explanation for the discrepancy between our results and those found by Aylward et al.^12^ is that this previous study included subjects with a mix of anxiety and depressive disorders, and a negative bias in learning may be more characteristic of depressive symptoms. Our PLS analysis provides some support for this speculation as we found that symptoms of depression were associated with elevated learning from threat, suggesting that such a bias in learning is associated more with depressive symptoms. It is also possible that the nature of our game-based online task engaged processes distinct from that of standard lab-based tasks. However, we believe this is unlikely since we replicate behavioural patterns shown in more traditional tasks, and also observed similar associations with anxiety in a previous lab-based study^15^. As such, we are confident that this is not simply due to the task used.

We found a similar pattern of enhanced learning from safety when examining a transdiagnostic factor representing compulsivity and intrusive thought. Although this factor has been shown to be associated with less model-based behaviour^6,25^, altered confidence judgements^26^, and action-confidence coupling^8^ in large-scale samples, to date it has not been investigated with regard to threat learning. Notably, we also found a weak relationship between this factor and uncertainty, whereby more compulsive individuals had higher certainty in their safety estimates, echoing previous work in perceptual decision making that showed this factor is associated with higher confidence estimates^8,26^. We only found weak relationships (where the posterior density estimate crossed zero) with the other two factors, representing anxious-depression and social withdrawal, in a direction indicative of lower safety probability estimates and higher uncertainty.

Perhaps our most striking results come from a data-driven approach where we derived components of psychopathology grounded in computational analyses of aversive learning behaviour. This method provides conceptually similar results to the factor analytic methods used in previous large-scale online studies^6^, but builds upon this work by incorporating behaviour into the process. Using PLS regression, we identified two latent components, one broadly associating greater learning from safety with physiological symptoms of anxiety and compulsivity, while the other associated greater learning from threat with depressive symptoms and social anxiety. Notably, this data-driven analysis also revealed relationships between aversive learning and impulsive behaviour, encompassing a symptom dimension that is typically studied in the context of reward processing^37^. Individuals scoring higher on these symptoms exhibited higher safety learning, which may explain previously observed relationships between impulsivity and risk tolerance^38^. Importantly, while this analysis was exploratory, we demonstrate its robustness through testing on held-out data, ensuring our results are not affected by overfitting^36^.

Overall, the present results add to the growing literature showing associations between psychopathology and learning under uncertainty. Previous studies using computational approaches have largely focused on learning about rewards and losses^10–12,27,39^, or perceptual learning^9^, and those that have used more aversive paradigms (using outcomes intended to evoke subjective anxiety), such as learning to predict electric shocks, have been limited by small samples^5,15,18,40^. As a result, the precise role played by aversive learning processes in psychiatric symptoms has been unclear. Our work adds to this literature by providing an account of how these processes relate to symptoms across a range of traditional diagnostic categories.

The results we report raise questions of importance for future work. In particular, the finding that more anxious individuals tend to overestimate safety likelihood runs counter to intuition, and further work is required to understand how this may relate to symptom expression. One speculative possibility is that a persistent underestimation of threat likelihood would lead to an abundance of aversive prediction errors, causing a state of subjective anxiety. An alternative explanation is the result reflects a tendency for highly anxious individuals to seek safety, and be resistant to leaving places associated with safety^41,42^. However, these hypotheses await direct testing, and it will be especially important to examine them in large-scale clinical samples, taking into account a broader range of psychiatric phenotypes.

A further important feature of this study is our development of a new online task for measuring aversive learning. A number of studies examining other aspects of learning and decision making in the context of psychiatric disorders have also availed of large samples recruited through online services^6,8,25,26^. However, it has been difficult to examine aversive learning in online environments, as aversive lab stimuli such as shock cannot be easily administered online. Only one study thus far has investigated threat-related decision making (although not learning) online, using monetary loss as an aversive stimulus^27^. A game-based design allowed us to design a task that required avoidance behaviour as well as evoke feelings of anxiety, taking advantage of the well-known ability of games to produce strong emotional reactions^43–47^, resulting in a paradigm which we believe provides a more valid assessment of aversive learning than more commonly used monetary-loss based tasks. Although qualitatively different from standard lab-based tasks, we observed similar patterns of biased learning to that seen in lab-based work^15^. An added benefit of our task is that it is highly engaging, and subjects reported feeling motivated to perform well. These features are not only important for the kind of large-scale online testing performed here. This task renders it feasible to measure aversive learning at regular intervals without subjects needing to physically visit the lab, a feature that could be of considerable utility in clinical trials.

One potential limitation of this study is a focus on a general population sample which, being recruited online, was not subject to the kind of detailed assessment possible offline. While this might limit applicability to clinical anxiety, other research indicates that findings from clinical samples replicate in samples recruited online^6,8^. Furthermore, it is increasingly recognised that clinical disorders lie on a continuum from health to disorder^48^. Although we did not deliberately set out to recruit individuals with clinically significant anxiety, 36% of our sample scored at or above a threshold designed for the detection of anxiety disorders on our measure of trait anxiety (see supplementary material). In light of this, and given limitations with research in clinical samples that includes medication load^49^ and recruitment challenges^50^, online samples provide an effective method for studying clinically-relevant phenomena. Additionally, it is important to note that the effects we observed were small, as in previous studies using large-scale online testing^6,25,25^. However, large samples provide accurate effect size estimates in contrast to the exaggerated effects that are common in studies using small samples^51^. Such small effects are unsurprising given the multifactorial nature of psychiatric disorders^52^. While we have shown aversive learning to be important, we acknowledge this is likely to be one of a multitude of processes involved in the development of these conditions.

In conclusion, our results demonstrate links between transdiagnostic symptoms of psychiatric disorders and mechanisms of threat learning and uncertainty estimation in aversive environments. The findings emphasise the importance of these processes not only in anxiety but indicate a likely relevance across a spectrum of psychopathology.

## Methods

### Ethics

This research was approved by the University College London research ethics committee (reference 9929/003). All participants provided informed consent and were compensated financially for their time at a rate of at least £6 per hour.

### Participants

We recruited 400 participants through Prolific^31^. Subjects were selected based on being aged 18-65 and having at least a 90% approval rate across studies they had previously participated in. As described in our preregistration, we used a precision-based stopping rule to determine our sample size, stopping at the point at which either the 95% highest posterior density interval (HPDI) for all effects in our regression model reached 0.15 (checking with each 50 subjects recruited) or we had recruited 400 subjects. The precision target was not reached, and so we stopped at 400 subjects.

### Avoidance learning task

Traditional lab-based threat learning tasks typically use aversive stimuli such as electric shocks as outcomes to be avoided. As it is not possible to use these stimuli online, we developed a game-based task in which subjects’ goal was to avoid negative outcomes. While no primary aversive stimuli were used, and subjects received no actual monetary reward, there is an extensive literature showing that video games without such outcomes evoke strong positive and negative emotional experiences ^43–47^, making this a promising method for designing an aversive learning task. In this game, participants were tasked with flying a spaceship through asteroid belts. Subjects were able to move the spaceship in the Y-axis alone, and this resulted in a one dimensional behavioural output. Crashing into asteroids diminished the spaceship’s integrity by 10%. The spaceship’s integrity slowly increased over the course of the task, however if enough asteroids were hit the integrity reduced to zero and the game finished. In this eventuality subjects were able to restart and continue where they left off. The overarching goal was to maximise the number of points scored, where the latter accumulated continuously for as long as the game was ongoing, and reset if the spaceship was destroyed. Subjects were shown the current integrity of the spaceship by a bar displayed in the corner of the screen, along with by a display of their current score.

Crucially, the location of safe spaces in the asteroid belts could be learned, and learning facilitated performance as it allowed correct positioning of the spaceship prior to observing the safe location. The task was designed such that without such pre-emptive positioning it was near impossible to successfully avoid the asteroids, thus encouraging subjects to learn the safest positions. Holes in the asteroids could appear either at the top or bottom of the screen (Figure 1A), and the probability of safety associated with either location varied independently over the course of the task. Thus, it was possible to learn the safety probability associated with each safety zone and adapt one’s behaviour accordingly. The probability of each zone being safe was largely independent from the other (so that observing safety in one zone did not necessarily indicate the other was dangerous), although at least one zone was always safe on each trial. Participants also completed a control task that required avoidance that was not dependent on learning, enabling us to control for general motor-related avoidance ability in further analyses (described in supplementary material). We elected a priori to exclude subjects with limited response variability (indicated by a standard deviation of their positions below 0.05) so as to remove subjects who did not move the spaceship. However, no subject met this exclusion criterion.

After completing the task, subjects were asked to provide ratings indicating how anxious the task made them feel and how motivated they were to avoid the asteroids, using visual analogue scales ranging from 0 to 100.

### Behavioural data extraction

For analysis, we treated each pass through an asteroid belt as a trial. Overall there were 269 trials in total. As a measure of behaviour, we extracted the mean Y position across the 1 second prior to observing the asteroid belt, representing where subjects were positioning themselves in preparation for the upcoming asteroid belt. This Y position was used for subsequent model fitting. On each trial, the outcome for each zone was regarded as “danger” if asteroids were observed (regardless of whether they were hit by the subject) or safety if a hole in the asteroid belt was observed.

### Computational modelling of behaviour

Our modelling approach focused on models that allowed the quantification of subjective uncertainty. To this end, we modelled behaviour using approximate Bayesian models that assume subjects estimate safety probability using a beta distribution. This approach is naturally suited to probability estimation tasks, as the beta distribution is bounded between zero and one, and provides a measure of uncertainty through the variance of the distribution. While certain reinforcement learning formulations can achieve similar uncertainty-dependent learning and quantification of uncertainty, we chose beta models as they have an advantage of being computationally simple. Empirically, these models have been used successfully in previous studies to capture value-based learning^53^, where they explain behaviour in aversive learning tasks better than commonly used reinforcement learning models^15,54^, a pertinent characteristic in the current task.

The basic premise underlying these models is that evidence for a given outcome is dependent on the number of times this outcome has occurred previously. For example, evidence for safety in a given location should then be highest when safety has been encountered many times in this location. This count can be represented by a parameter *A*, which is then incremented by a given amount every time safety is encountered. Danger is represented by a complementary parameter *B*. The balance between these parameters provides an indication of which outcome is most likely. Meanwhile, the overall number of outcomes counted influences the variance of the distribution and hence the uncertainty about this estimate. Thus, uncertainty is highest when few outcomes have been observed. The exact amount by which *A* and *B* are updated after every observed outcome can be estimated as a free parameter (here termed *τ*), and we can build asymmetry in learning into the model, so that learning about safety and danger have different rates, allowing updates for *A* and *B* to take on different values (here termed *τ*^*+*^ and *τ*^*-*^).

Such a model is appropriate in stationary environments, when the probability of a given outcome is assumed to be constant throughout the experiment. However, in our task the probability of safety varied, and so it was necessary to build a forgetting process into the model. This is achieved by incorporating a decay (represented by parameter *λ*) which diminishes the current values of *A* and *B* on every trial. The result of this process is akin to reducing the number of times they have been observed, and maintains the model’s ability to update in response to incoming evidence. It would also be possible to build asymmetry into the model here, where subjects could forget about positive and negative outcomes at different rate. However, testing this model in pilot data revealed that separate decay rates for each valence were not recoverable. Estimates for *A* and *B* are therefore updated on each trial (*t*) according to the following equation for both safety zones independently (termed *X* and *Y* here). Both zones are updated on every trial, as subjects saw the outcome associated with both simultaneously. This formed the basis of all the probabilistic models tested:

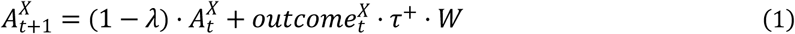

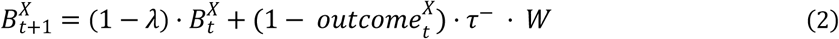

We also observed in pilot data that subjects tended to be influenced more by outcomes occurring in the zone they had previously chosen, an effect likely due to attention. On this basis, we incorporated a weighting parameter that allowed the outcome of the unchosen option to be down-weighted by an amount shown in the above equation (*W*) determined by an additional free parameter, *ω*.

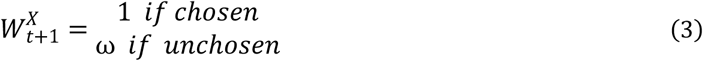

We can calculate the estimated safety probability for each zone (*P*) by taking the mean of this distribution:

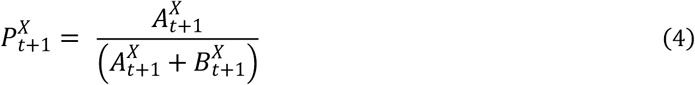

Similarly, we can derive a measure of uncertainty on each trial by taking the variance of this distribution.

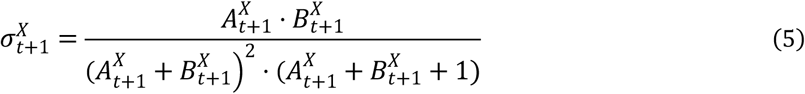

In order to fit our model to the observed behaviour, we require an output that represents the position of the spaceship on the screen. This position (*pos*) was calculated based on the safety probability of the two safety zones, such that the position was biased towards the safest location and was nearer the centre of the screen when it was unclear which position was safest.

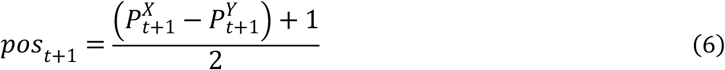

Further models elaborated on this basic premise, and full details are provided in supplementary material. For completeness, we also tested two reinforcement learning models, a Rescorla-Wagner model and a variant of this model with different learning rates for better and worse than expected outcomes^32^, both of which are described in supplementary material. However, we focus on the probabilistic models due to their ability to represent uncertainty naturally; our primary aim was not to differentiate between probabilistic and reinforcement learning models, but to use previously validated models to provide insights into the relationship between aversive learning, uncertainty, and psychopathology.

Models were fit with a hierarchical Bayesian approach using variational inference implemented in PyMC3, through maximising the likelihood of the data given a reparametrised beta distribution with a mean provided by the model and a single free variance parameter. Model fit was assessed using the Watanabe-Akaike Information Criterion (WAIC)^55^, an index of model fit designed for Bayesian models that accounts for model complexity. Parameter distributions were visualised using raincloud plots^56^.

### Measures of psychiatric symptoms

Our first set of hypotheses focused on state/trait anxiety and intolerance of uncertainty. These were measured using the State Trait Inventory of Cognitive and Somatic Anxiety (STICSA)^57^ and the Intolerance of Uncertainty Scale (IUS)^58^ respectively. We also wished to examine how behaviour in our task related to the three transdiagnostic factors identified by Gillan et al. (2016), based on factor analysis of a range of psychiatric measures. To measure these factors more efficiently, we developed a reduced set of questions that provided an accurate approximation of the true factor scores, details of which are provided in supplementary material.

### Regression models

Bayesian regression models were used to investigate relationships between behaviour and psychiatric measures, predicting each behavioural measure of interest from the psychiatric measures. Our dependent variables were parameters and quantities derived from our model, which represented the way in which an individual learns about safety probability and how they estimate uncertainty. Specifically, we used the two update parameters from our model (*τ*^*+*^ and *τ*^*-*^, referring to the extent to which subjects update in response to safety and danger respectively) and the mean safety probability and uncertainty estimates across the task (generated by simulating data from the model with each subject’s estimated parameter values). Crucially, the fact that task outcomes were identical for every subject ensured these values were dependent only on the manner by which subjects learned about safety, not the task itself.

These models were constructed using Bambi^59^ and fit using Markov chain Monte Carlo (MCMC) sampling, each with 8000 samples, 2000 of which were used for burn-in. All models included age and sex as covariates, along with performance on our control task to account for non-learning related avoidance ability. For analyses predicting state and trait anxiety and intolerance of uncertainty, we constructed a separate model for each variable due to the high collinearity between these measures. For analyses including the three transdiagnostic factors, these were entered into a single model. When reporting regression coefficients, we report the mean of the posterior distribution along with the 95% highest posterior density interval (HPDI), representing the points between which 95% of the posterior distribution’s density lies. All analyses were specified in our preregistration. We did not correct for multiple comparisons in these analyses as our approach uses Bayesian parameter estimation, rather than frequentist null hypothesis significance testing, and as such multiple comparison correction is unnecessary and incompatible with this method^60^.

### Partial least squares regression

To provide a data-driven characterisation of the relationship between task behaviour and psychiatric symptoms, and identify transdiagnostic components that are grounded in both self-report and behaviour, we used partial least squares (PLS) regression to identify dimensions of covariance between individual questions and the measures derived from our modelling. We excluded the STICSA state subscale from this analysis, so that only trait measures were included. To ensure robustness of these results, we split our data into training and testing sets, made up of 75% and 25% of the data respectively. To identify the appropriate number of components within the training set, we used a 10-fold cross-validation procedure, fitting the model on 90% of the training data and evaluating its performance on the left-out 10%. The mean squared error of the model’s predictions was then averaged across test folds to provide an index of the model’s predictive accuracy with different numbers of components, using cross-validation to reduce the risk of overfitting

Once the number of components was determined, we validated the model’s predictions by testing its predictive accuracy on the held-out 25% of the data. To provide a measure of statistical significance we used permutation testing, fitting the model on the training data 1000 times with shuffled outcome variables and then testing each fitted model on the held-out data, to assess its predictive accuracy when fitted on data where no relationship exists between the predictors and outcomes. This procedure provides a null distribution, from which we can then determine the likelihood of observing predictive accuracy at least as high as that found in the true data under the null hypothesis.

Recent work has highlighted the risks inherent in PLS-like methods when used in high dimensional datasets^36^, namely that they can easily be overfit resulting in solutions that do not generalise beyond the data used to fit the model. Our approach avoids these problems by evaluating the performance on our model 25% of the data that has been held out from the model fitting stage.

### Preregistration and data availability

The main hypotheses and methods of this study were preregistered on the Open Science Framework (https://osf.io/jp5qn). The data-driven PLS regression analysis was exploratory. Data is available at https://osf.io/b95w2/ and code is available at https://github.com/tobywise/online-aversive-learning.

## Acknowledgements

T.W. is supported by a Wellcome Trust Sir Henry Wellcome Fellowship (206460/17/Z). R.J.D. holds a Wellcome Trust Investigator award (098362/Z/12/Z). The Max Planck UCL Centre is a joint initiative supported by UCL and the Max Planck Society. The Wellcome Centre for Human Neuroimaging is supported by core funding from the Wellcome Trust (203147/Z/16/Z). We thank Evan Russek for comments on an earlier version of this manuscript.

