## Supplementary material for "Associations between aversive learning processes and transdiagnostic psychiatric symptoms revealed by large-scale phenotyping"

### Supplementary materials

#### Control task

Participants completed a control task that required avoidance that was not dependent on learning, in order to quantify each subjects' avoidance ability resulting from non-learning related factors. This allowed us to control for general motor-related avoidance ability in further analyses and ensured that relationships between our behavioural variables and psychopathology were not simply a result of non-learning factors such as reaction time. This task was similar to the to the main task, however a group of asteroids was always positioned to appear at the same Y position as the subject's current location. As in the main task, subjects had to avoid asteroids, but this was dependent only on the ability to move out of the way of oncoming asteroids rather than the ability to learn their position.

#### Computational models

We tested a range of computational models of the behaviour on our task. We focused on probabilistic models, termed "asymmetric leaky beta" models due to the fact that they update in response to safety and danger asymmetrically, and incorporate a leak parameter to imbue them with the flexibility to update estimates in response to incoming information rather than assuming safety probabilities are fixed. The basic machinery of this model family, along with the full best fitting model, is described in the main text, however here we describe variations that we also tested in addition to the reinforcement learning models tested.

##### Asymmetric leaky beta

This is the most basic model, which the other models build upon. This is described in the main text.

##### Softmax-transformed asymmetric leaky beta

This model builds upon the basic model by incorporating a softmax transform on the estimated optimal position, as described in the main text.

Additionally, based on pilot data we had noted that subjects tended to position themselves nearer to either the top or bottom of the screen, and tended to avoid the centre. To encourage similar performance from our model, a softmax function with inverse temperature ($\beta$) as a free parameter was applied to the estimated position:

$$pos_{t+1} = \frac{e^{(\beta \cdot pos_{t+1})}}{e^{(\beta \cdot pos_{t+1})} + e^{(\beta \cdot (1 - pos_{t+1}))}} \quad (1)$$

Although softmax functions are typically used for converting continuous value estimates to choice probabilities in decision making tasks, we emphasise that here this method is used simply to produce position estimates that avoid the centre of the screen rather than the conventional usage of computing choice probability.

#### Asymmetric leaky beta incorporating choice stickiness

Our best fitting model included a "stickiness" parameter, which caused the chosen safety zone to be more likely to be chosen on the following trial (other models tested and compared with the winning model are described fully in supplementary material). This was achieved by boosting the value of the chosen option (determined by whether the position was above or below the centre) by raising the estimated safety probability of the option to the power of S, a free parameter bounded between zero and one governing the degree of stickiness. Here an exponential effect was chosen over an additive one as it ensures the probability does not take a value above 1.

$$P_{t+1}^{X} = \begin{matrix} {P_{t+1}^{X}}^{S} \ if \ chosen \\ P_{t+1}^{X} \ if \ unchosen \end{matrix} \quad (2)$$

#### Variance weighted asymmetric leaky beta

This model weights the two screen locations based on their variance when calculating the optimal Y position. The location with the lowest variance is weighed most highly, as follows. Firstly, a variance bias measure was calculated, representing the ratio between the variance of the top and bottom zones (labelled X and Y here).

$$\sigma_t^{bias} = \frac{\sigma_t^X}{\sigma_t^X + \sigma_t^Y} \quad (3)$$

This bias measure was then used to weight the safety probability estimates of the two options, such that the probability estimate was highest for the option with the lowest variance. This weighting was itself dependent on a free modulatory parameter $\pi$, to allow the amount of variance weighting to differ on an individual subject basis.

$$Pw_t^X = P_t^X \cdot (1 - \sigma_t^{bias} \cdot \pi) \quad (4)$$

$$Pw_t^Y = P_t^Y \cdot (1 - (1 - \sigma_{bias})_t \cdot \pi) \quad (5)$$

#### Upper confidence bound asymmetric leaky beta

This model also incorporates uncertainty into the position calculation, through an upper confidence bound rule. This means that rather than using the mean of the distribution as the location's safety estimate, an upper part of the distribution is used, the exact level of which is estimated as free parameter $\omega$.

$$UCB_t^X = P_t^X + \omega \cdot \sqrt{\sigma_t^X} \tag{6}$$

The position is then calculated as follows:

$$pos_t = \frac{\left(UCB_t^X - UCB_t^Y\right) + 1}{2} \tag{7}$$

#### Rescorla-Wagner

We also tested two reinforcement learning models for comparison. The first of these was a standard Rescorla-Wagner model (1) which updates safety estimates based on prediction errors ($\delta$) weighted by a learning rate ($\alpha$), which is estimated as a free parameter.

$$P_{\mathrm{t+1}}^{\mathrm{X}} = P_{\mathrm{t}}^{\mathrm{X}} + \alpha \cdot \left(\mathrm{outcome}^{\mathrm{X}} - P_{\mathrm{t}}^{\mathrm{X}}\right) \tag{8}$$

Probability estimates are combined in the same way as the beta models and are also then passed through a softmax function with a free inverse temperature parameter to decrease the likelihood of positions near the centre of the screen.

#### Dual learning rate Rescorla-Wagner

The second reinforcement learning model we tested was an extension of the Rescorla-Wagner model (2) that allowed better and worse than expected outcomes to have differential effects on safety updates by introducing a separate learning rate for each

$$\delta = \mathrm{outcome}^{\mathrm{X}} - \mathrm{V}_{\mathrm{t}}^{\mathrm{X}} \tag{9}$$

$$P_{\mathrm{t+1}}^{\mathrm{X}} = P_{\mathrm{t}}^{\mathrm{X}} + \begin{matrix} \alpha^{+} \cdot \delta \text{ if } \delta_t > 0 \\ \alpha^{-} \cdot \delta \text{ if } \delta_t < 0 \end{matrix} \tag{10}$$

Probability estimates are converted to a single position estimate as in the previous models.

#### Task validation

First, any task that seeks to measure aversive learning should evoke a feeling of subjective anxiety in a majority of subjects. To assess this  subjects rated how anxious the task made them feel on a scale ranging from 0-100. The mean rating provided was 40.89 (SD = 30.45), although visual inspection of their responses indicated a strongly bimodal distribution (Figure 2A, main text). Notably, 76.5% of subjects reported a rating above 10 (indicating at least some anxiety). Based on this pattern we conclude  the task did evoke a feeling of subjective anxiety in a majority of subjects.

Second, the subjective effects produced by the task should relate to real-world individual differences; a task that successfully invokes subjective anxiety should do so to a greater extent in subjects who are generally more anxious. To test this, we assessed whether task-induced anxiety was predicted by state and trait anxiety. Spearman correlations revealed significant positive relationships with both these measures (state $\rho$=0.25, p<.001; trait $\rho$=0.22, p<.001), indicating that individuals reporting more general anxiety also found the task more anxiety-inducing.

Finally, the task should produce behaviour that resembles that of existing tasks. One task behavioural characteristic frequently observed in learning from positive and negative outcomes is a tendency to learn faster from outcomes that are negative than those that are positive[12,15,32]. To provide a simple behavioural measure of subjects' avoidance tendencies, we calculated the degree to which they moved their position following safe and dangerous outcomes at their chosen location. As expected, this revealed subjects tended to shift their position more following danger than following safety outcomes, indicating they adapted their behaviour in the task, and learnt about the safest locations, as in standard avoidance learning tasks (Figure 2E, main text).

#### Model validation

Given the centrality of this model in our analyses, we sought to verify the validity of its parameters and the data generated from the model. If valid, these should be clearly related to model-free behaviour and task outcomes, and so we began by testing this. First, due to an interest in asymmetric learning about safety and punishment, the two parameters of the preferred model with greatest relevance to our hypotheses were update rates for safety and danger. To verify that the estimated parameters truly reflected behavioural tendencies, we examined correlations between their estimated values and the tendency to stay following safety and move following danger across subjects (Figure 3D). These were robustly correlated, showing that subjects moved more following danger than safety, and providing confidence that these parameters related to relevant behavioural measures.

Second, we verified that values generated by simulating data from our model using best-fitting parameters were related to model-free behavioural measures. Here we focused on two model-derived values that were central to our subsequent analyses, namely uncertainty and safety value. We validated these measures by examining their relationships with the extent to which subjects changed their position on each trial. If our uncertainty measure is valid, subjects should change their position to a greater extent when they are more uncertain. Similarly, if our safety value measure is valid then subjects should change their position more when there is a smaller difference in value between the two zones (indicating that one is not clearly better than the other). We tested this using a hierarchical Bayesian regression model predicting change magnitude from uncertainty (averaging across the two zones) and value difference between zones within subject. As expected, this showed positive effect of uncertainty ($\beta$=0.15, 95% highest posterior density interval (HPDI)=0.14, 0.16), indicating that subjects moved more when uncertain, and a negative effect of value difference ($\beta$=-0.04, 95% HPDI=0.04, 0.03), indicating that subjects moved more when there was a less clear difference between the value of the two options (Figure 3C). This provides confidence that these model-derived measures are indeed representing uncertainty and safety value.

Finally, as an additional validation step, we examined how our modelling results compared to those obtained from computational models of traditional lab-based tasks[12,15,32]. We compared update rates for danger and safety in our task, expecting to see that subjects update faster in response to danger than safety, as shown in previous work[15]. This revealed a negative bias in the values of these parameters whereby subjects tended to update to a greater extent in response to danger than safety ($t(400) = 26.76$, $p < .001$, Figure 3E), in line with prior studies. It could be argued that this a result of subjects being forced to move when encountering danger, and can stay when encountering safety. However, three aspects of the task argue against this. First, subjects do not have time to move substantially when they observe danger (this makes learning and prediction essential). Second, a one-off dangerous outcome does not necessarily mean the zone is now the least safe, and subjects should often return if they did move following danger. Last, behaviour is dependent on the outcomes of *both* safety zones; a safe outcome in the alternative zone is as informative as a dangerous one in the current zone. Overall, the presence of such biased learning provides confidence that the task detects broad behavioural patterns seen in existing tasks.

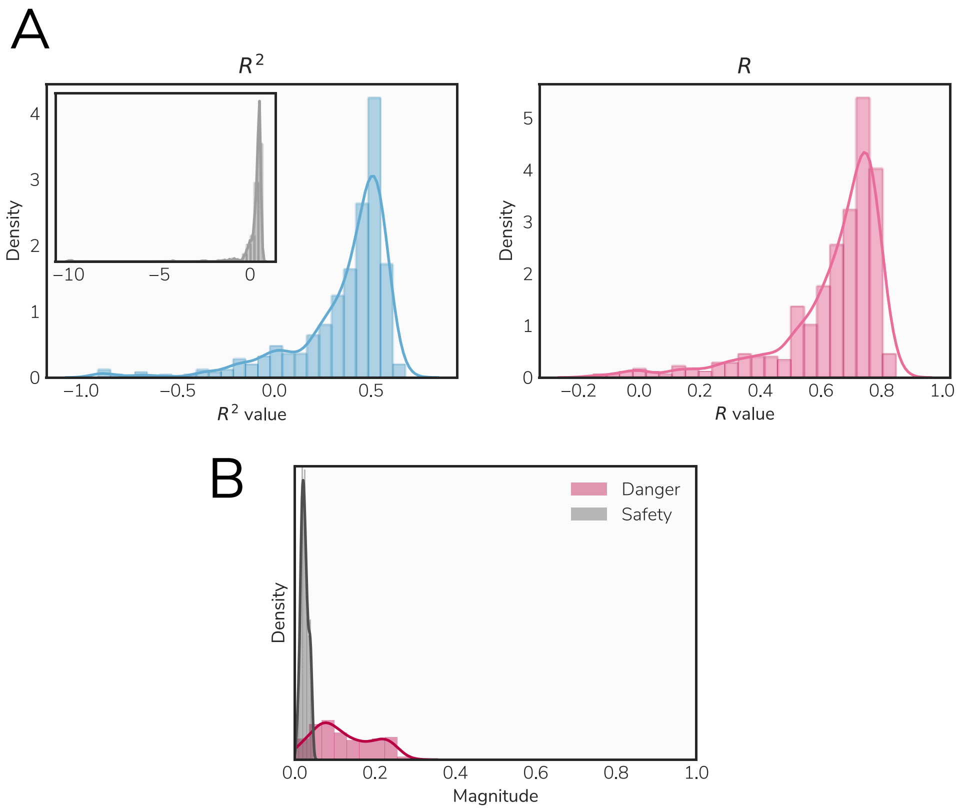


Figure S1. Correspondence between simulated and true data. A) Distributions of $R^2$ and $R$ values across subjects, showing a high correspondence between the real and simulated data. The inset plot in the first panel shows the full distribution of $R^2$ scores (which can take any value below or equal to 1), including a small number of subjects with poor model fit, while the main figure shows the values between -1 and 1. B) Magnitude of position changes following dangerous and safe outcomes in simulated data, showing that simulated subjects tend to change position to a greater extent in response to danger than safety, as seen in data from real subjects.

#### Correspondence between real and model-generated data

While our winning model provided a better fit to the data than other candidate models, it is important to ensure that our model truly produced data that was similar to subjects' real behaviour. We checked this in two ways: First, we calculated the $R^2$ value and Pearson's correlation between the true data and data simulated from our model with the estimated parameters for each subject, to ensure that we did indeed achieve a high correspondence between model-generated and true data. As these data were heavily skewed, due a small number of subjects where the model did not provide a good fit to the data, we report the median and interquartile range of the scores across subjects. These were 0.44 (0.26) and 0.70 (0.16) for $R^2$ and Pearson's $R$ respectively (Figure S1A), indicating good concordance between model-generated and true data.

Second, we checked that a basic pattern of behaviour that emerged in the true data was also present in the model-generated data. Subjects tended to change their position more following a dangerous outcome than a safe outcome, as would be expected if they are learning to avoid threat (as shown in Figure 2A). We repeated this analysis on our simulated data, and observed the same pattern of results (Figure S1B).

#### Parameter recovery

We parameter recovery tests on our winning model to ensure that we were able to accurately estimate parameter values. This involved simulating data from the model using 500 combinations of parameter values, each randomly drawn from gaussian distributions with a mean of 5 and standard deviation of 0.2 (setting any values generated above 1 to 1 and below zero to zero). We then fit our winning model to this simulated behaviour, and finally compared the recovered parameter estimates to those used to generate the data. This revealed generally high correlations between parameter sets. Most importantly, recovered $\tau^p$ and $\tau^n$ were correlated with the values used to generate the data at $R$ = 0.68 and 0.61 respectively, indicating that we were able to recover their values with reasonable accuracy. Full results are shown in Figure S2.

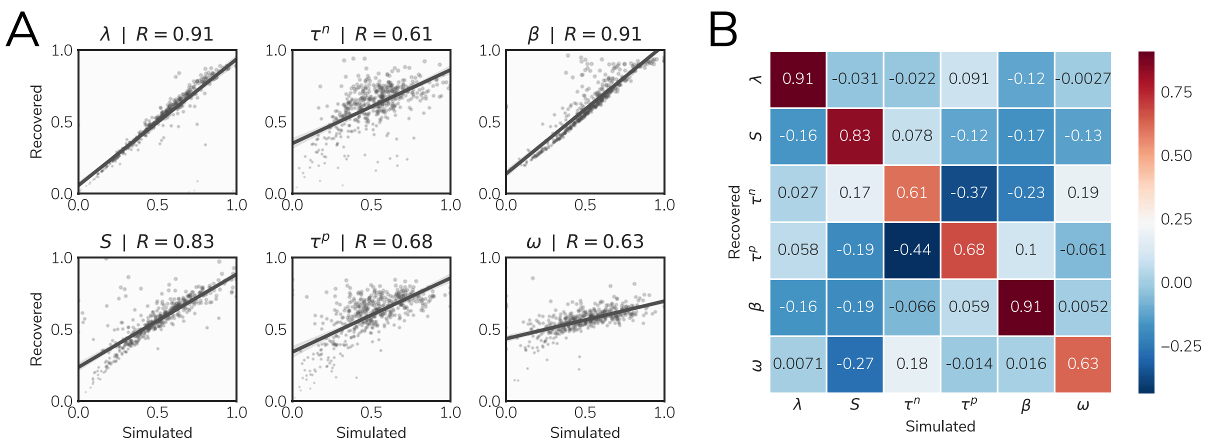


Figure S2. Parameter recovery results. A) Correlations between the value used to generate data and the recovered parameter value across all parameters individually. B) Correlation matrix showing Pearson R correlations between generating values and recovered values for all combinations of parameters.

#### A reduced set of questions for measuring transdiagnostic factors

We wished to investigate relationships with three transdiagnostic factors developed by Gillan et al., (2016). However, to reduce the number of questions used to determine scores on these three factors for each subject, and hence the time taken to complete the task, we used a data-driven approach to select the most important questions for determining factor scores. To achieve this, we used lasso regularised regression to predict each subject's factor score from responses to individual questions in data from the study by Rouault et al (4). Performing this analysis with a range of values of the hyperparameter C, which governs the degree of regularisation, produced a model that included varying numbers of questions as predictors. The ability of these models to predict the true factor scores was assessed using five-fold cross validation, whereby the model was trained on 80% of the data and tested on the remaining 20%, with this procedure repeated across combinations of training and test data and the prediction $R^2$ averaged across these five folds. Plotting this across values of C, and numbers of retained questions, allowed us to select a point at which we were able to achieve satisfactory accuracy with an acceptable number of questions. This resulted in a set of 63 retained questions out of an initial 225 (Figure S3A, Table S1), which resulted in $R^2$ values of .91, .81, and .89 for the three factors (Figure S3B).

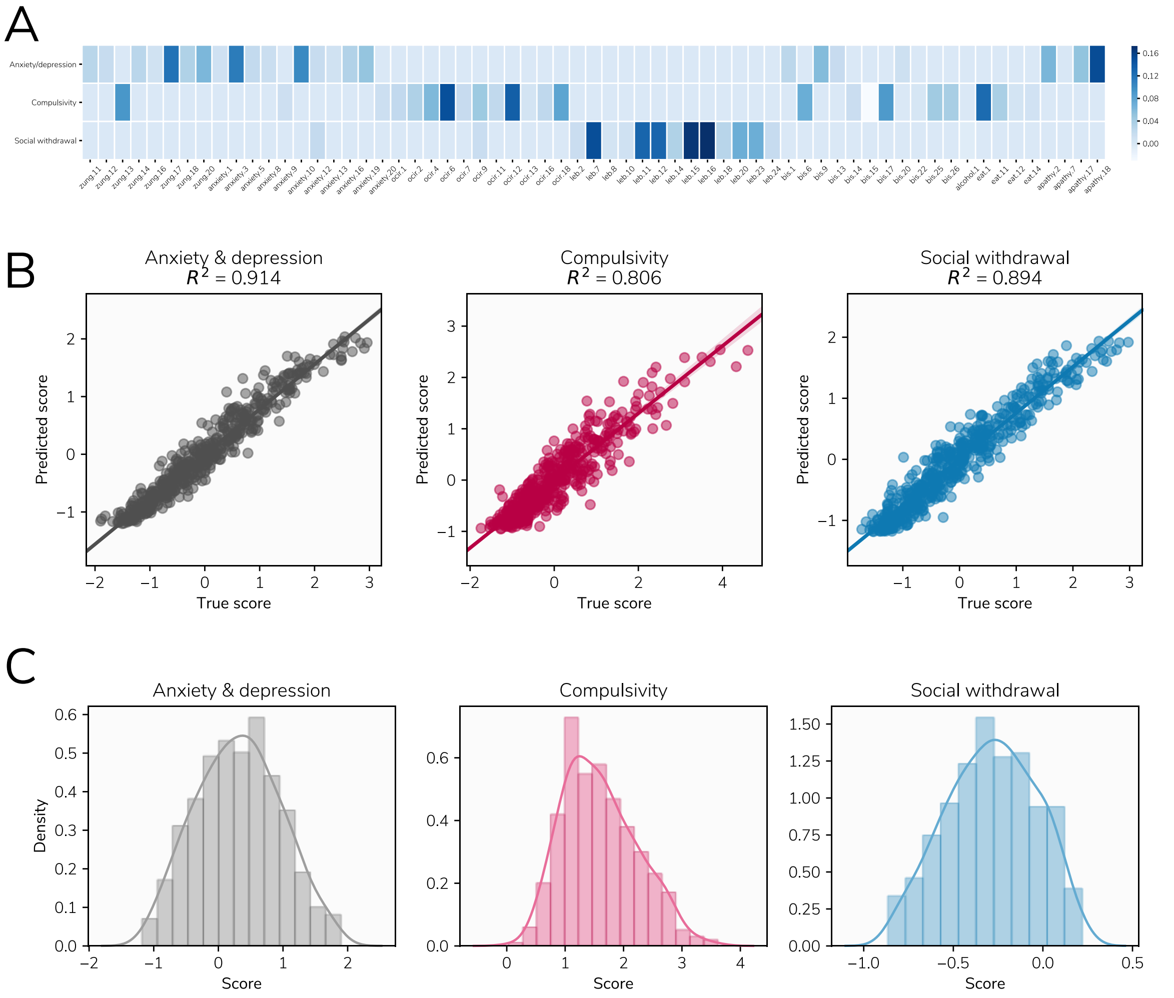


Figure S3. Result of question reduction procedure, finding a reduced set of questions that allow prediction of subject scores on the three transdiagnostic factors identified by Gillan et al. (2016). A) Weights of the retained questions in the regularised logistic regression model, demonstrating which questions are predictive of each factor. B) True factor scores from the study by Rouault et al. (2018) plotted against the cross-validated predicted factor scores from our model, demonstrating a high degree of accuracy. C) Distributions of factor scores in the data from the current study.

Table S1. Questions included in the reduced set of items used to approximate scores on the three factors derived by Gillan et al., 2016 (5).

| Measure | Item numbers |
|---|---|
| Zung Depression Scale (5) | 11, 12, 13, 14, 16, 17, 18, 20 |
| State Trait Anxiety Inventory (trait subscale) (6) | 1, 3, 5, 8, 9, 10, 12, 13, 16, 20 |
| Obsessive Compulsive Inventory (Revised) (7) | 1, 2, 4, 6, 7, 9, 11, 12, 13, 16, 18, |
| Liebowitz Social Anxiety Scale (8) | 2, 7, 8, 10, 11, 12, 14, 15, 16, 18, 20, 23, 24 |
| Barratt Impulsivity Scale (9) | 1, 6, 9, 13, 14, 15, 17, 20, 22, 25, 26 |
| Alcohol Use Disorder Identification Test (10) | 1 |
| Eating Attitudes Test (11) | 1, 11, 12, 14 |
| Apathy Evaluation Scale (12) | 2, 7, 17, 18 |

#### Effects of task motivation

Subjects reported their motivation to avoid asteroids in the task on a visual analogue scale, ranging from 0 to 100. It is possible that task motivation may be affected by psychopathology, and to explore this possibility we constructed a regression model predicting task motivation from state and trait anxiety, intolerance of uncertainty, and the three transdiagnostic factors used in the main analysis. The strongest effect in this analysis was that of intolerance of uncertainty, whereby subjects scoring higher on this measure rated their motivation higher (mean $\beta$ = 0.15, 95% HPDI = 0.02, 0.28). Full results are shown in Figure S4.

Following this, we repeated our main analyses with task motivation as a covariate to ensure that our results were not better explained by effects of motivation than psychopathology per se. Results remained almost unchanged from the original analyses (Figure S5).

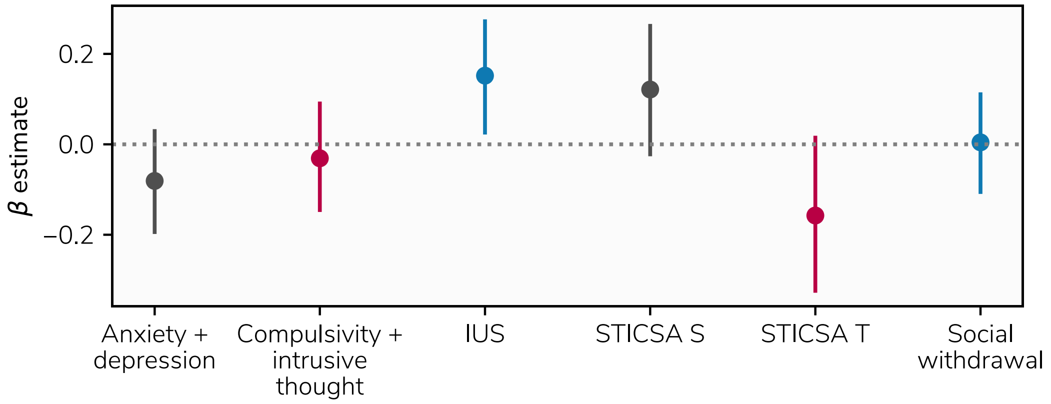


Figure S4. Effects of psychopathology on reported task motivation.

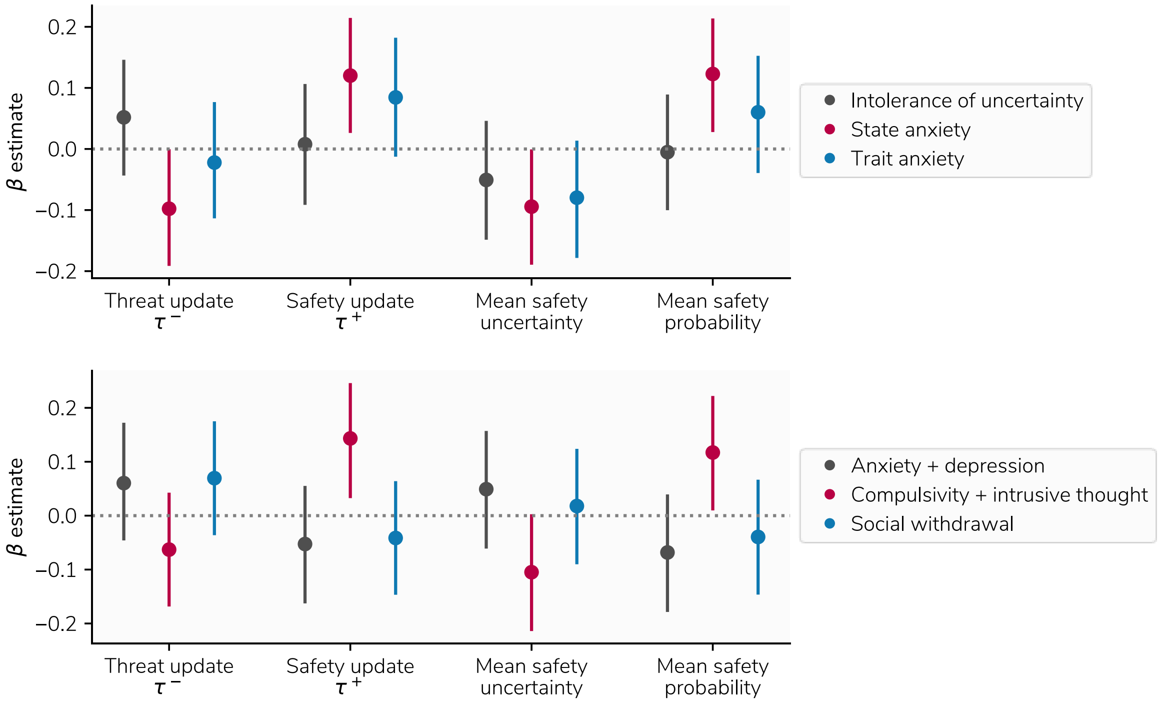


Figure S5. Relationships between model-derived behavioural measures and psychopathology, controlling for task motivation.

#### Subjects with clinical levels of anxiety

Although we did not aim to recruit subjects with clinically-significant symptoms of anxiety, given the high prevalence of anxiety disorders it would not be unexpected to find subjects with such levels of anxiety in a large general population sample. We did not use any measure designed to diagnose anxiety disorders, and as such is impossible to determine for certain how many subjects would meet diagnostic criteria. However, it is possible to use approximate thresholds on other measures to provide an indication of the proportion of the sample who may have clinically significant anxiety symptoms. For our measure of trait anxiety, the STICSA

(6), such a threshold has been identified by van Dam et al., (7). This study indicated that a score of 43 on the scale was able to distinguish individuals diagnosed with anxiety disorders from healthy controls with 74% accuracy. In our data, 144 of our 400 subjects (36%) scored above this threshold (Figure S3), indicating that a substantial proportion of our sample are likely to be experiencing clinically significant symptoms of anxiety.

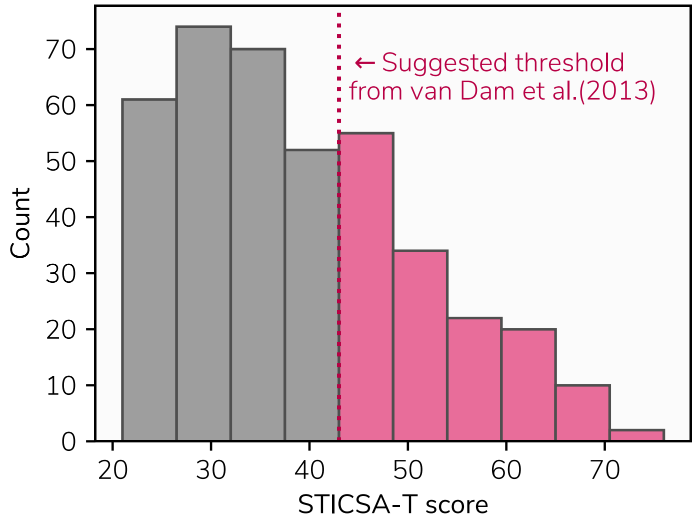


Figure S6. Proportion of subjects scoring above a threshold indicating likely presence of a clinical anxiety disorder determined by van Dam et al. (7). The dotted line indicates the threshold of 43 on the STICSA trait scale, while the histogram bars coloured pink represent subjects scoring above this threshold.

#### Analyses using an approximate binary anxiety status indicator

We next sought to investigate whether the aforementioned anxiety disorder status, based on the threshold identified by van Dam et al, could be used to predict behavioural measures in the same way as our continuous indexes of psychopathology. To test his, we labelled the 144 subjects identified above as likely anxiety disorders, and the 144 lowest scoring subjects on the STICSA-T as likely healthy. We then used this as a binary predictor, along with sex and age as covariates, in regression models with our model-derived behavioural measures as dependent variables. The results of this analysis showed only weak effects, as seen in Figure S7. This may be a due to a number of factors. First, this is a very approximate method for identifying anxiety disorders, and so our groups are unlikely to fully conform to true anxiety disorders and truly healthy. Second, this approach resulted in reduced numbers of subjects overall, reducing statistical power. Last, such a binary measure contains less information than the continuous measure it is derived from.

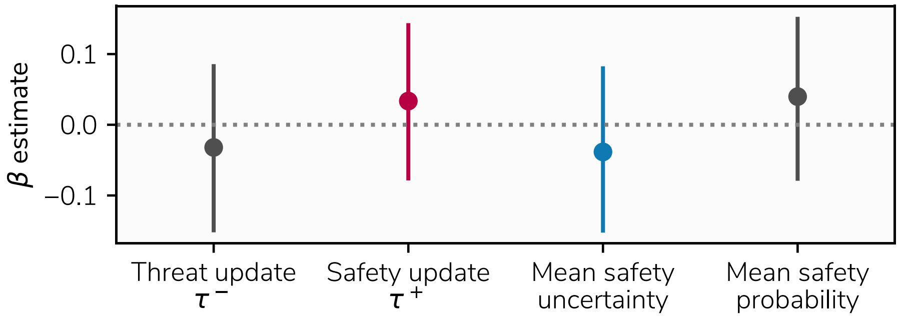


Figure S7. Effects of binary approximate anxiety disorder status on model-derived behavioural measures.

### Full item loadings for PLS analysis

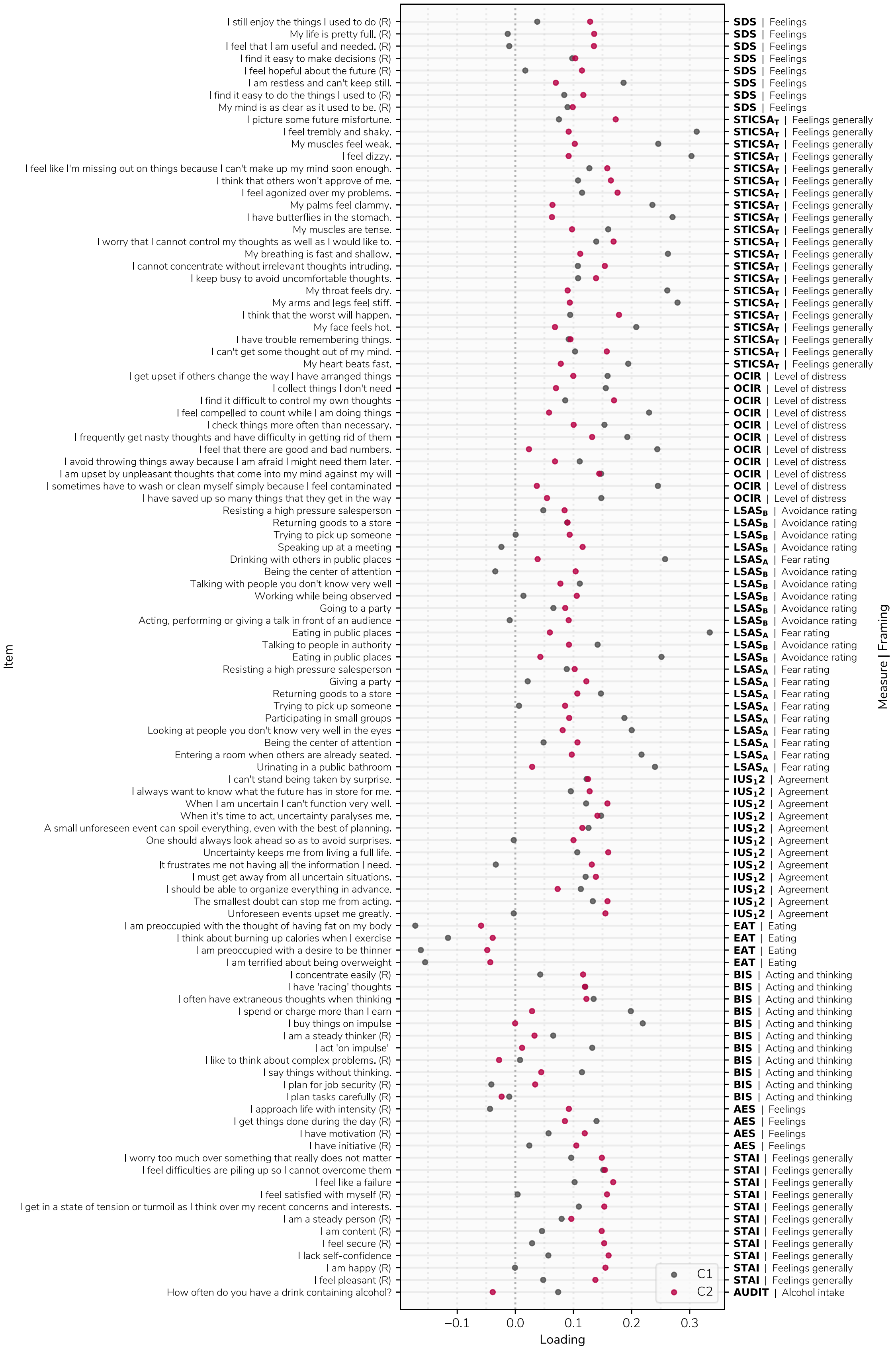


Figure S8. Full item loadings for the PLS analysis on both components.
